# Electrical properties, accuracy, and multi-day performance of gelatine phantoms for electrophysiology

**DOI:** 10.1101/2020.05.30.125070

**Authors:** Amani Yousef Owda, Alexander J. Casson

## Abstract

Gelatine based phantoms for electrophysiology are becoming widely used as they allow the controlled validation of new electrode and new instrumentation designs. The phantoms mimic the electrical properties of the human body and allow a pre-recorded electrophysiology signal to be *played-out*, giving a known signal for the novel electrode or instrumentation to collect. Such controlled testing is not possible with on-person experiments where the signal to be recorded is intrinsically unknown. However, despite the rising interest in gelatine based phantoms there is relatively little public information about their electrical properties and accuracy, how these vary with phantom formulation, and across both time and frequency. This paper investigates ten different phantom configurations, characterising the impedance path through the phantom and comparing this impedance path to both previously reported electrical models of Ag/AgCl electrodes placed on skin and to a model made from ex vivo porcine skin. This article shows how the electrical properties of the phantoms can be tuned using different concentrations of gelatine and of sodium chloride (NaCl) added to the mixture, and how these properties vary over the course of seven days for a.c. frequencies in the range 20–1000 Hz. The results demonstrate that gelatine phantoms can accurately mimic the frequency response properties of the body–electrode system to allow for the controlled testing of new electrode and instrumentation designs.

## I. Introduction

Electrophysiological sensing, measuring the electrical activity of the brain (electroencephalography, EEG), heart (electrocardiography, ECG), muscles (electromyography, EMG), and eyes (electrooculography, EOG), is an extremely common technique used in both clinical practice and fundamental research [1]. While electrophysiological sensing instrumentation is well developed, there are significant ongoing research efforts into improving the sensing performance [2]. This is in terms of both the hardware, making it smaller, more robust to artefacts and easier to use (for example [3], [4]), and the electrodes, with a wide range of electrode materials and approaches investigated in recent years (for example [5], [6]). These new electrodes particularly have the objective of not requiring a conductive gel to lower the body contact impedance [7].

While patient simulators are available for testing instrumentation (for example the well known Fluke Medsim [8]), these simulate only the physiological waveform and not the full body– electrode–instrumentation connection and so do not allow the complete electrode–instrumentation system to be validated. Instead, for testing new electrodes and complete systems early stage human or animal work is typically performed. However, this introduces ethical or animal license considerations, and is labour intensive, both of which limit the rapid, iterative, and large scale screening of different electrode approaches. Moreover, when using a human or animal model the true electrophysiological signal present is not known, which makes the validation of systems very challenging as no ground truth is available to compare against [2].

In contrast, body phantoms permit the controlled evaluation of electrodes and instrumentation over a wide range of realistic parameters such as noise, motion, and artefacts [9], and are commonly used in fields such as Magnetic Resonance Imaging (MRI). While physical phantoms have their own limitations compared to computational modelling approaches, physical phantoms offer the benefits of incorporating environmental noise, real world interference, and electrode or instrumentation non-idealities to allow realistic early stage testing. They also do not require a realistic model of the novel electrodes and the contact they make with the body. However, historically, electrophysiological phantoms have been extremely simple, with melons widely used [10], [11]. This is because melons have similar electrical properties to the human head [12], and the size of a melon is roughly the same size as a head [11].

Clearly, much improved phantoms are possible and these are now starting to emerge, particularly for EEG verification, Fig. 1. Generally these more advanced head phantoms can be categorised into three types: single layer (or homogeneous) phantoms; multi-layer phantoms (usually three layers representing scalp, skull, and brain); and real-tissue phantoms that use animal tissue or cadavers to have actual biological tissue as part of the phantom.

**Fig. 1.**
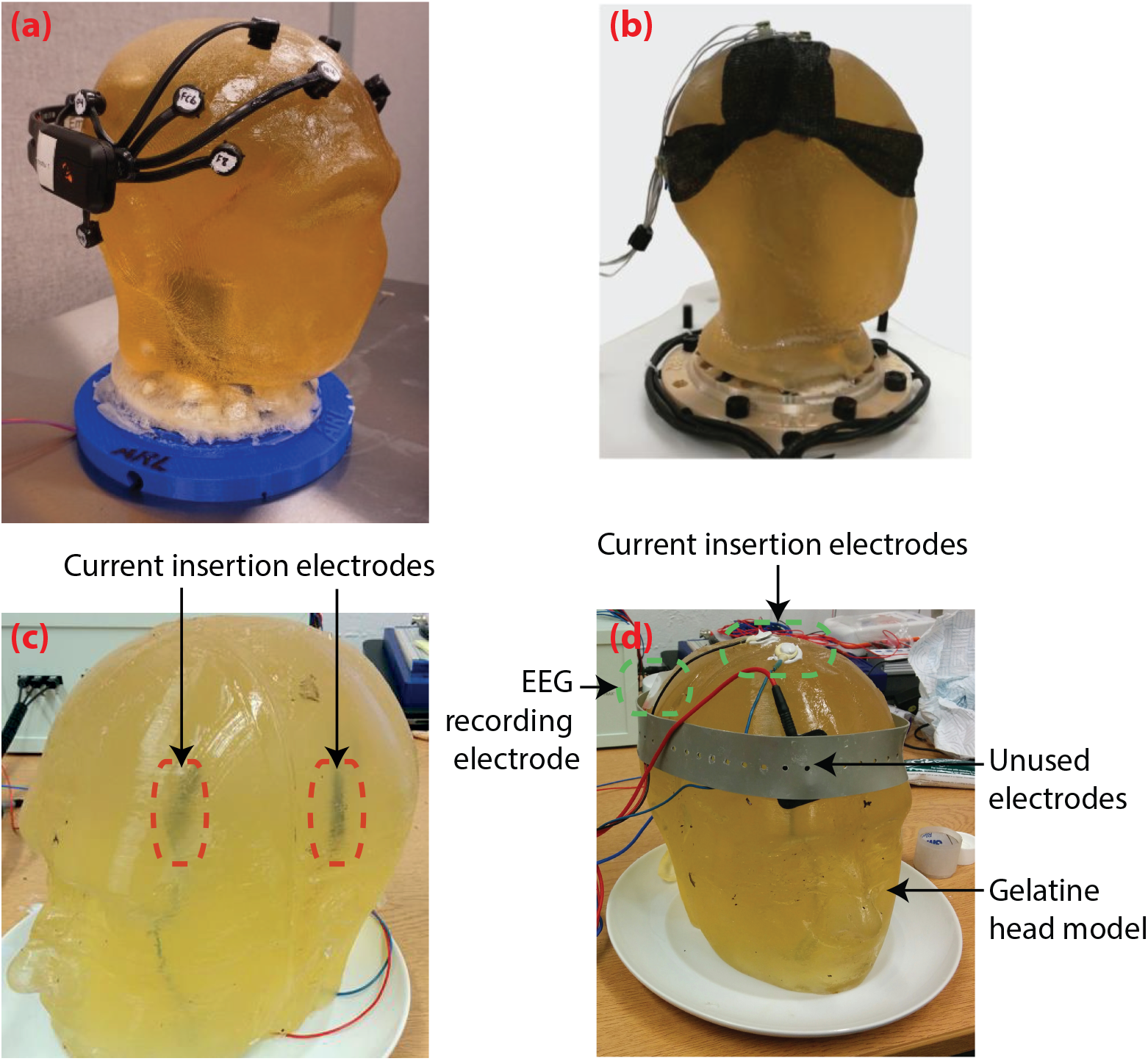
Examples of current homogeneous gelatine electrophysiology phantoms. (a) A head phantom from the US Army Research Laboratory with manufacturing details available on the open access web site [14]. Image taken from the public domain [20]. (b) Gelatine head phantom from [15] which was placed on a robotic platform to simulate movements due to walking. Reprinted under the CC-BY license from [15]. (c) Head phantom from our lab with two internal electrodes for replaying a pre-recorded signal which can be collected by electrodes on the phantom surface. Reprinted by permission from Springer Nature: Biomedical Engineering Letters, [2], © 2019. (d) Head phantom from our lab for EEG combined with brain stimulation (tES electrodes labelled as *unused* in this picture). © 2017 IEEE. Reprinted, with permission, from [21].

Single layer phantoms have received the most attention, and all of the phantoms illustrated in Fig. 1 are made of a single homogeneous layer. These are typically based upon ballistics grade gelatine, and aim to mimic the electrical properties of the body with electrical sources embedded within the gelatine to allow a pre-recorded electrophysiological signal to be *played-out* and recorded again on the surface of the phantom using the test electrodes or instrumentation. They therefore provide the test electrodes or instrumentation with a known signal to record and compare against. For example, [13] used homogeneous gelatine phantoms, tuning the contact impedance profile by using different concentrations of sodium chloride (NaCl). The methods and materials for making this phantom have been released in an open source manner [14], and this phantom has also been used to investigate EEG components collected during walking motions [15] where the true EEG is normally unknown as it is obscured by artefacts. Cuboid shaped phantoms were used in [5] to verify the performance of 3D printed EEG electrodes, and to demonstrate how the electrode performance varied with contact force. Some research groups have even applied transcranial Electrical Stimulation (tES) to these phantoms to investigate the tES artefact removal problem from EEG where a known EEG signal can now be provided [16], [17]. [18] suggested agar as an alternative to gelatine for the recording of slow EEG potentials, while recently [19] made use of an agarose gel swollen with a saline solution.

More sophisticated, multi-layer phantoms, again particularly for heads have also been proposed. For brain stimulation situations [22] presented a three layer model for comparing the distribution of brain stimulation currents to a finite element model. For electrical impedance tomography [23] presented a multi-layer 3D printed head phantom, while [24] used a saline water phantom for the same application. Focusing on electrophysiology multi-layer works are more limited, with [25] presenting a three-layered model consisting of a brain (made of urethane resin), skull (made of silicone), and scalp (made of silicone) for testing EEG caps and indicated that realistic scalp electric potentials were generated. [26] used Tx-151 gel [27], deionised water, sodium chloride, agar and sucrose for investigating EEG and ECG source imaging. This phantom was shown to be stable for use over the course of 8 hours.

The result is that a wide number of different materials have been investigated for making electrophysiology body phantoms (gelatine [2], [5], [13], [15], [16], wax [28], silicon [28], plastic clay [29], and plastic moulds [22]), with gelatine being by far the most common. Based on this, gelatine has also been used as the base material for real-tissue phantoms. [30] used a phantom made of a dry real skull, filled with a conductive medium (solidified saline gelatine) to validate EEG source localisation. [31] used a human head skull phantom filled with gelatine to investigate the effect of dipole localisation on EEG and magnetoencephalogram (MEG).

Despite this significant interest, works characterising the electrical properties of gelatine and other materials for mimicking the body are limited. [13] is a one-page investigation on tuning the electrical properties of gelatine phantoms over a 5 Hz to 1.5 kHZ range using different concentrations of NaCl, but there is no information on how this compares to biological tissues, or how the electrical properties change over time. [17], [19] match only the d.c. conductivity and do not consider the a.c. frequency response. [32] investigated the a.c. properties of gelatine and agar phantoms over the range 100–500 Hz with varying levels of NaCl, but again did not compare these to biological tissues, or how the electrical properties change over time. [33] investigated the static and dynamic mechanical properties of gelatine phantoms having 10%, 20% and 30% of gelatine concentrations. Thermal and mechanical properties of gelatine phantoms were investigated in [34], with milk added to control these. As a result, it is clear that gelatine based phantoms are being widely used by the research community, but with very limited open information on their electrical properties, representativeness, and how to control these properties.

This paper presents a wide ranging investigation into the performance of gelatine phantoms for electrophysiology. We investigate the electrical impedance of phantom devices made with both different concentrations of gelatine and different concentrations of NaCl for obtaining control over the electrical properties. We pay particular attention not only to the impedance across a.c. frequencies, but also to the group delay present, allowing accurate transmission of information through the phantom without introducing timing distortions. In addition, we measure how these factors change over a week of storage/use to demonstrate phantom stability, reusability, and repeatability. We then compare the accuracy of the gelatine phantoms to both an electrical equivalence model of the human body which has been reported previously in the literature [35], and to measurements of ex vivo porcine skin. The result is a demonstration of the controllability of gelatine phantom properties across a.c. frequencies, and a demonstration that they can be customised to be highly accurate models of human tissues with usability over the course of a week. To our knowledge this is the first direct comparison of these factors for gelatine with animal tissues and electrical equivalent models.

Section II overviews the electrical properties of the human body, and particularly the head, that gelatine phantoms need to mimic, together with established electrical models of these. Section III then overviews our procedure for the creation of gelatine phantoms. Methods and results for establishing the electrical properties of these are given in Section IV and Section V respectively, together with a comparison of these factors to electrical models and to an ex vivo porcine skin model. Finally, the results are discussed in Section VI and conclusions given in Section VII.

## II. Biological target to model

There are many different factors of interest for including in a phantom model, accurately mimicking electrical properties, mechanical properties and physical properties (e.g. hair, sweat glands) of the human body. Extensive work on MRI phantoms has investigated mimicking the mechanical [9], [28], [36], [37] and physical properties [34], [38]. For electrophysiological phantoms, and for this work, of most importance is having a good electrical equivalent of the human body and its tissues.

The electrical properties of the human body have been well characterised previously [35]. For the head (which has multiple layers of brain, skull, skin, and electrode interface) the equivalent electrical model is shown in Fig. 2 for different types of electrodes. In all cases the equivalent circuit model is similar, varying depending on the number of resistor–capacitor tanks present, and the component values.

**Fig. 2.**
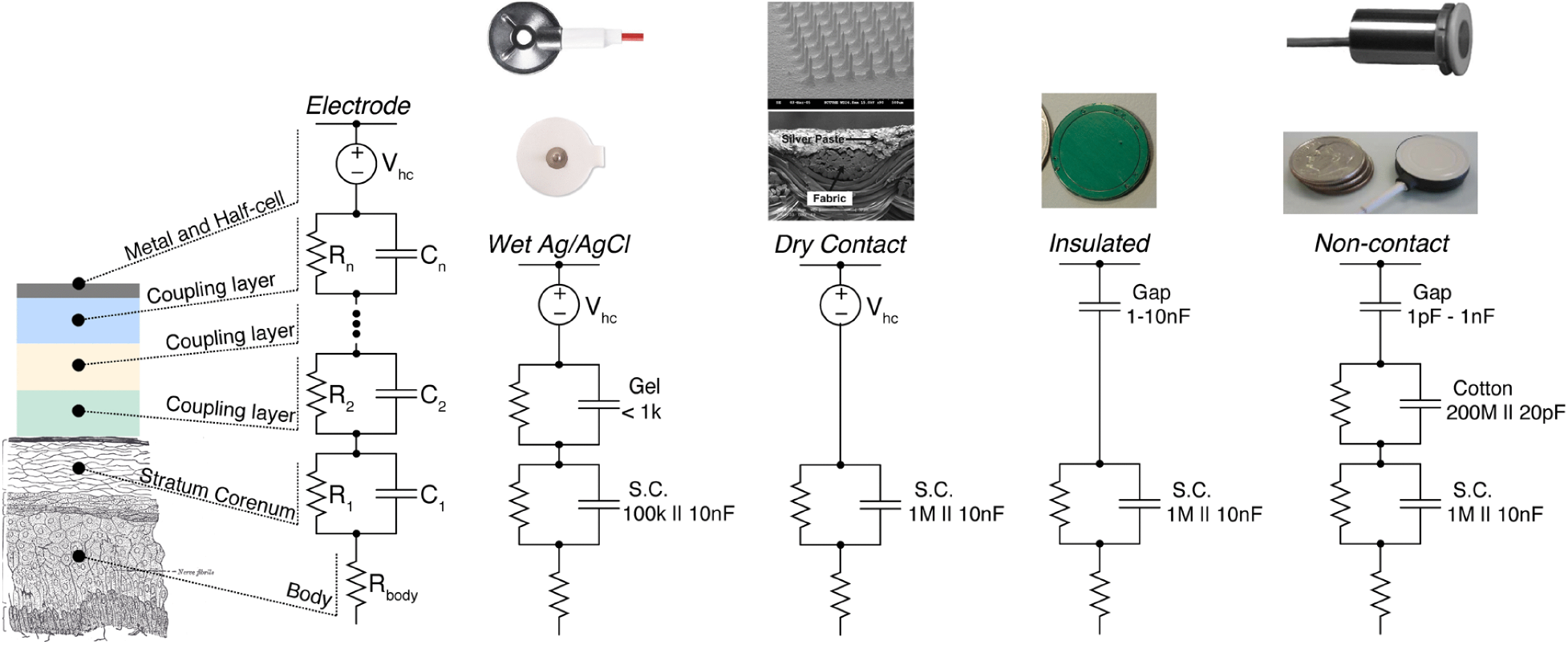
Different types of electrophysiological recording electrodes and the associated equivalent circuits; this includes wet-contact Ag/AgCl electrodes which are the model focused on in this work. © 2010 IEEE. Reprinted, with permission, from [35].

In this work we focus on modelling the wet Ag/AgCl system from Fig. 2. This consists of a bulk resistance (value unspecified) representing the body and internal tissues, and a 100 kΩ resistor and 10 nF capacitor representing the stratum corneum and giving a tank with a pole break point at 159 Hz (given by 1/2*πRC*). A second resistor–capacitor tank then represents the electrode coupling giving the system a second pole, with component values not specified. To estimate the break point of this tank we performed measurements of conductive gel using a widely used commercial gel (ABRALYT HiCl EEG electrode gel, Easycap, DE). The conductive gel was placed on a flat plate and spread to a uniform 2 mm thickness. A digital multimeter was then used for measuring the resistance and capacitance with probes placed 2 cm apart. Across multiple readings the resistance was found to be in the range 200–300 Ω and the capacitance in the range 60–120 *μ*F, agreeing with the <1 kΩ impedance given in Fig. 2 at low frequencies. Based on these measurements there is a second pole break point to model in the frequency range 5.3–15.9 Hz. In the wet Ag/AgCl system from Fig. 2 two zeros are also present, for which the locations can be worked out analytically. (The resulting equations are not compact and so not given here.)

From Fig. 2 the body to electrode contact impedance (*Z*) is a complex quantity that consists of a real part (*Z*_*real*_) and an imaginary part (*Z*_*imag*_)

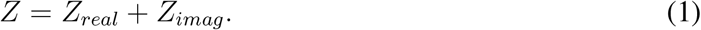

From this the magnitude (*Z*_*mag*_) and phase of the contact impedance can be calculated as

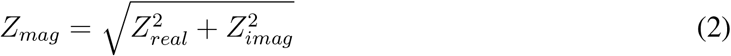

and

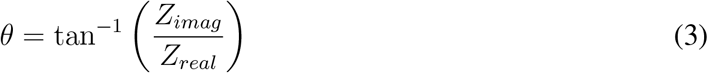

respectively. Here *Z*_*mag*_ is in Ω, and is commonly measured when setting up electrodes to ensure that a good body contact is present. For example, clinical standards for EEG ask for the contact impedance to be below 5 kΩ for a good contact to be present [1] (although with modern instrumentation good recordings can often be obtained with much higher impedances present [1]).

The phase response of electrodes is less commonly reported, but is important for electrode design and characterisation as a non-linear phase response will introduce time domain distortion into the collected signal. The group delay (*τ*) of the electrical connection is given by

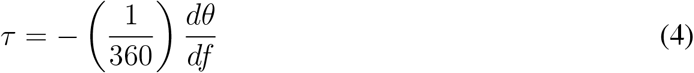

in seconds, assuming that *θ* is in degrees and *f* is in Hz. With non-linear phase responses the group delay will not be constant, and will vary with frequency such that some frequency components pass through the collection system more quickly than others, leading to distortion. Typical group delays are in the *μ*s range and so small compared to most electrophysiological signal time-spans [5] but not necessarily negligible. Measurements of group delay are included in our phantom characterisation results presented here.

## III. Test phantom creation

Electrophysiology phantoms were created using ballistics gelatine (ClassiKool, 240 blooms) that is cost effective (approximately *£*17 per kilogram) [39], conductive [33], easy to build and shape, easy to handle and use, portable, accessible and available with no restrictions. The gelatine powder was mixed with deionised water using different percentages of mass and then heated in a standard microwave for a period of time of 1 minute. This period was chosen as it is sufficient to dissolve the gelatine completely in the water without generating air bubbles unlike other studies [13], [14] where boiling water and a deforming agent were used to generate the phantom and remove the bubbles. A decision was made to use a microwave approach in this study since the phantom can be removed from the microwave at any time, before the generation of bubbles, giving more control over bubble formation.

After heating, the mixed gelatine was placed into a cuboid container (65×61×21 mm) to make the phantom. While we have previously moulded the phantom into realistic body shapes (e.g. a head in [21]) this was mainly for aesthetic purposes and to provide space for connecting multiple electrodes. Our more recent phantoms have used a cuboid shape [5] as this provides a uniform test structure with a flat surface for electrode connection, and our cuboids are sized to fit in with test infrastructure which allows fixed forces to be applied to electrodes to measure the effect of pressure and similar (as reported in [5]). This cuboid shape is now our preferred mould for day-to-day electrode testing where 1–2 electrode configurations/types are considered at a time, and is used in this paper rather than a full head shape. Our cuboid container was 3D printed using a flexible filament (NinjaFlex Semiflex) to allow easy removal once set. (For head shaped phantoms we similarly 3D print a mould, using a head shaped template available at [40].)

For providing an electrical source inside the phantom a standard Ag/AgCl electrode (Easycap, DE) was placed inside the gelatine mixture before it solidifed as a reference electrode to measure the contact impedance. Most electrophysiology phantoms have multiple electrodes inside [2], [14] so they can develop a potential difference (either directly or via a current injection) which can be measured on the phantom surface as the electrophysiological signal. For this study we included only one electrode inside the gelatine as the aim is to characterise the electrical properties of the phantoms, i.e. to measure the impedance path between the electrode inside the phantom and an electrode on its surface. The reference electrode was placed in the middle of the phantoms a few minutes after the mixture was obtained from the microwave. Since the mixture is getting thicker and stickier over time this method permitted controlled placement of the electrode inside the phantoms.

After the placement of the internal electrode the mixture was covered and placed in a lab fridge for 24 hours at a temperature of 2.3 °C to be used the next day (counted as day 1). An example of a complete cuboid phantom is shown in Fig. 3.

**Fig. 3.**
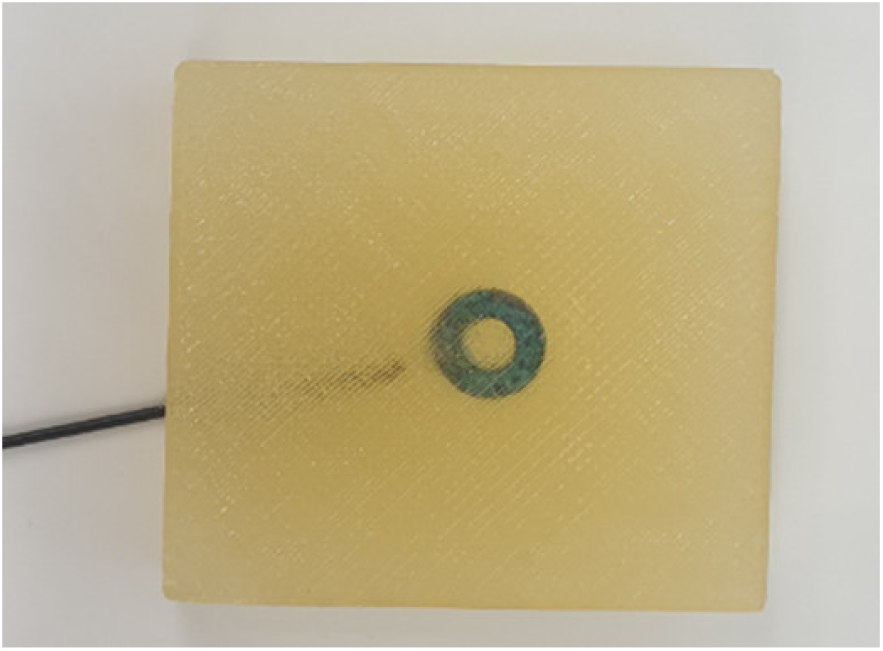
A cuboid gelatine phantom with a conventional Ag/AgCl electrode inside to act as a signal source.

To control and explore the electrical properties of the gelatine phantom the gelatine concentration used per mass of water was varied, and varying amounts of table salt NaCl was dissolved in the water before the gelatine powder was added. Ten different configurations are explored in this work:

i. 16.9% of gelatine concentration per mass, no added NaCl.
ii. 21.7% of gelatine concentration per mass, no added NaCl.
iii. 26.7% of gelatine concentration per mass, no added NaCl.
iv. 31.6% of gelatine concentration per mass, no added NaCl.
v. 30.0% of gelatine concentration per mass, no added NaCl.
vi. 30.0% of gelatine concentration per mass, and 0.5% of NaCl concentration per mass.
vii. 30.0% of gelatine concentration per mass, and 1% of NaCl concentration per mass.
viii. 30.0% of gelatine concentration per mass, and 2% of NaCl concentration per mass.
ix. 30.0% of gelatine concentration per mass, and 5% of NaCl concentration per mass.
x. 30.0% of gelatine concentration per mass, and 10% of NaCl concentration per mass.

The range of NaCl concentrations used matches that from the previous literature [13], and allows different body tissues to be mimicked. The gelatine values are non-round as a percentage due to the fixed volume of the phantom and the percentages of both the water and gelatine changing as more gelatine is added.

## IV. Experimental methods

### A. Impedance measurements

Impedance measurements were carried out using an Agilent 4284A precision LCR meter as shown in Fig. 4. A gold standard Ag/AgCl electrode was connected to the phantom and the magnitude and phase of the impedance path between the internal and external Ag/AgCl electrodes measured. The external electrode was held in place with the mechanical testing structure described in [5] with a 135 g mass used to fix the contact pressure between the electrode and the gelatine phantom. This surface Ag/AgCl electrode is common to all of the measurements carried out and included in the reported impedances. Impedance measurements were obtained at 20 Hz (the minimum frequency of the LCR meter) and then over the frequency band from 50 Hz to 1000 Hz in a regular step size of 50 Hz.

**Fig. 4.**
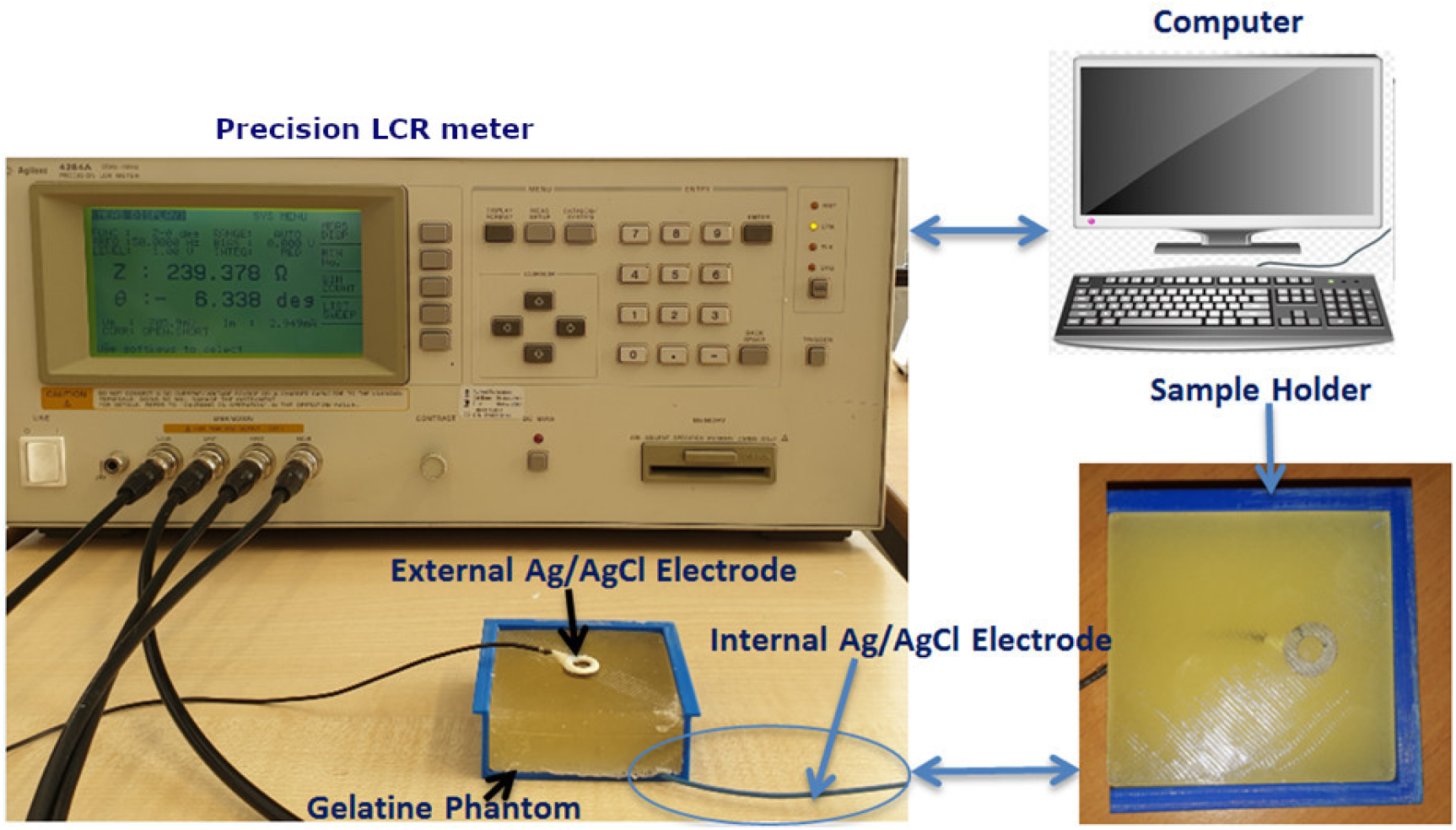
Precision LCR meter set up used for measuring the impedance of the gelatine phantoms in combination with a standard Ag/AgCl electrode.

Contact impedance measurements were performed directly after the phantoms were removed from the fridge at a temperature of 2.3±1 °C (measured using a thermometer (Hanna Instruments, HI935002)). The contact impedance responses of the gelatine phantoms were measured within a period of time of less than 5 minutes to maintain a constant temperature on the phantom during the experimental measurements as the electrical properties of the phantom vary with temperature [29], [41]. After each use the phantom was placed in a plastic box and returned to the fridge to be used the next day. These measurements were repeated at the same time of each day (10 am) over a period of time of one week excluding weekend days i.e. (day 1 to day 4 and then day 7) where all phantoms were made on a Monday.

### B. Electrical model fitting

Data from the precision LCR meter was exported to MATLAB and used to fit to the model given in Fig. 2 for the wet Ag/AgCl model, using the curve fitting toolbox and a least squares optimisation procedure. This focused on fitting the two poles present in this system to show how the gelatine phantom–electrode system matches the frequency response of the human body–electrode system. Different fits were calculated for each characterisation performed (each phantom on each day) with the optimisation procedure constrained to select similar values to the previous day’s fit. The breakpoints of the resistor and capacitor tanks were first fitted based upon the collected phase response. These breakpoints were then used as the starting estimate to fit the magnitude response and select precise values for the resistor and capacitor which would not *jump* between days. (The same break frequency could be obtained from multiple resistor–capacitor combinations.)

Fitted lines are included on all of the reported impedance curves in Section V, with the accuracy of these discussed specifically in Section V-E. Values on the fitted lines are plotted at 0.1 Hz intervals, not only at phantom frequency measurement points. As the full equation for the resistor–capacitor model from Fig. 2 is available no extrapolation is needed for plotting these points.

### C. Ex vivo porcine skin fitting

To validate the modelling compared to a biological target an ex vivo porcine skin model was also used. Previous works have shown this is a close mimic of human tissues [42], [43], but not compared it to gelatine phantoms. Porcine skin samples used in this research were purchased from the food chain via a commercial butchers shop. The work was logged with our faculty governance and no animal license was required.

Ex vivo porcine skin samples were taken from the back region of a healthy animal having age of eight months and an average weight of 70 kg. This region is free from hair follicles [44] which presents a limitation for both porcine skin and gelatine phantoms. The purchased sample of porcine skin was cut to have a rectangular shape and dimensions (140×60 mm, 10 mm thickness). The contact impedance measurement set up is shown in Fig. 5 with the porcine skin sample placed in the middle of the two electrodes. Apart from the electrode placement, all impedance measurements were carried out identically to those for the gelatine phantoms.

**Fig. 5.**
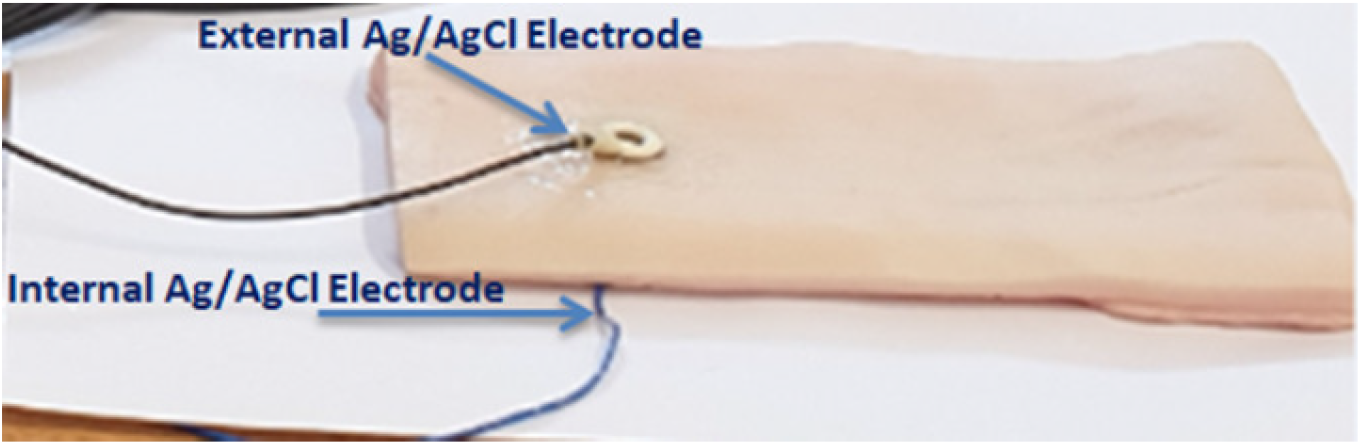
Illustration of electrode placement for ex vivo porcine skin impedance measurements.

## V. Results

### A. Effect of gelatine concentration

Fig. 6 shows the contact impedance response (magnitude and phase) of phantoms i–iv, having concentrations of gelatine 16.9–31.6%, and how this changes over the course of seven days. The results show that the magnitude response obtained from these phantoms increases over time and falls with frequency, and the phase response becomes less negative over both time and frequency. These trends are due to the water content that is evaporated (or absorbed) over time making the phantom drier [45]. The contact impedance response results obtained from the electrical model (discussed in more detail in Section V-E) were found to fit well with the experimental measurements of the gelatine phantoms. This indicates that the base gelatine phantom is an accurate physical model of the skin–electrode system that is to be represented. The gelatine is intrinsically conductive and its concentration can be used to control the properties of the phantom.

**Fig. 6.**
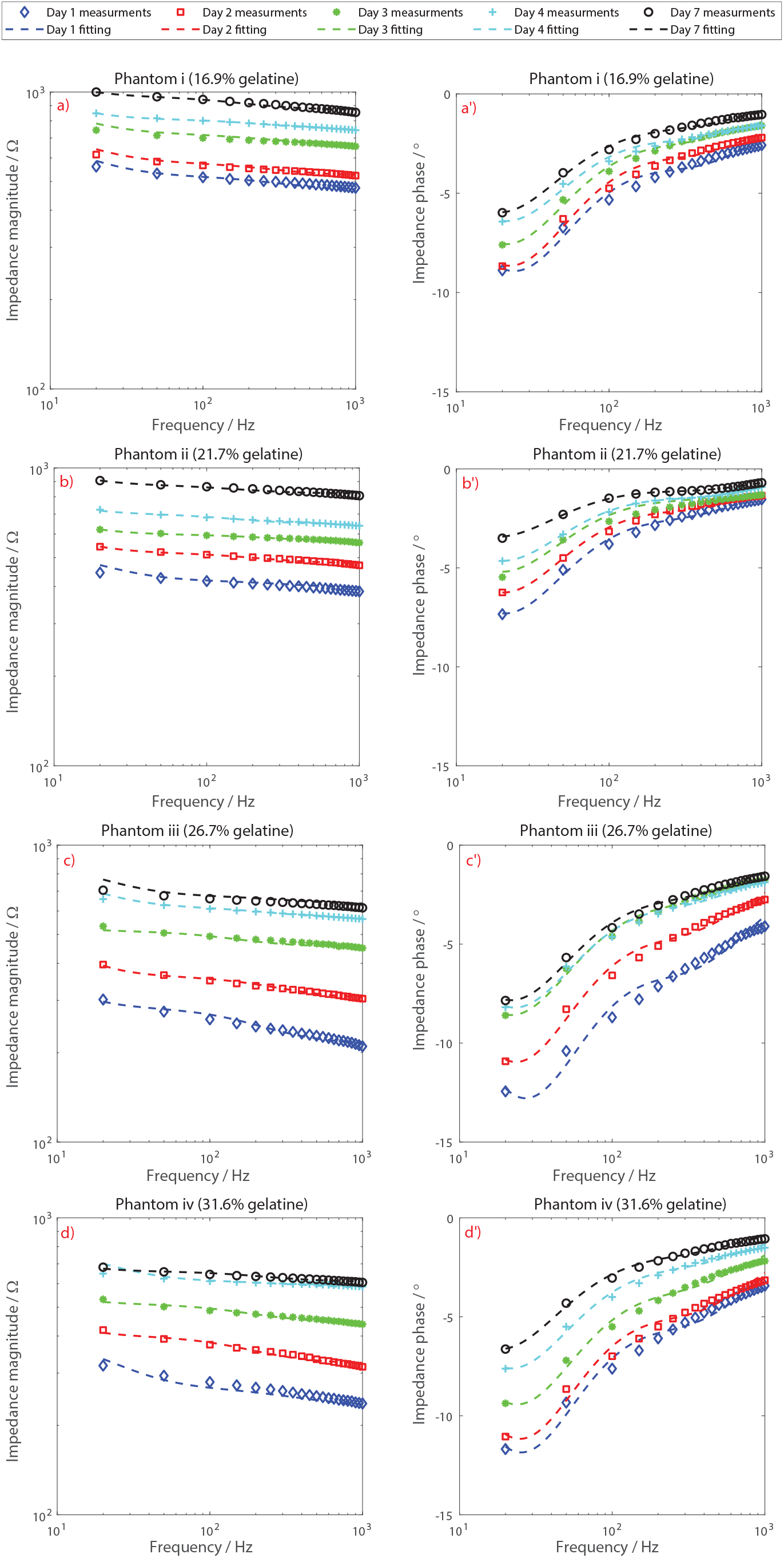
Impedance magnitude and phase responses for gelatine phantoms i–iv (in panels a–d respectively) with concentrations of gelatine 16.9–31.6%, and how these vary over the course of 7 days. Dashed lines indicate the fit of the second order electrical model from Fig. 2.

A comparison between phantoms having different concentrations of gelatine in Fig. 6 shows that the contact impedance response of phantoms having 26.7% and 31.6% of gelatine concentrations per mass is lower than those having gelatine concentrations of 16.9% and 21.7%. This indicates that increasing the gelatine concentration increases the conductivity (*σ*) of the phantoms. The minimum magnitude of the contact impedance was 216 Ω with a mean phase response of −4.1° achieved at 1000 Hz in day 1 for phantom iii. In contrast the maximum magnitude of the contact impedance does not exceed 1 kΩ with a mean phase response of −5.96°, achieved at 20 Hz in day 7 for phantom i. Without hair present to make the connection to the skin difficult low contact impedances are possible in all cases, matching those predicted from Fig. 2. The range of impedances from 216–1000 Ω is small compared to the 5 kΩ contact impedance accepted for clinical use. Although the impedances increase over time, staying below 5 kΩ means that the phantoms are durable over the course of 7 days, and a further increase in impedance (i.e. use longer than 7 days) would still give physiological accurate impedance magnitude values.

The corresponding group delays are shown in Fig. 7. The measurements indicate that the group delay varies over time, and from phantom to phantom, although this effect is not large with all of the readings at the same frequency point being within 100 *μ*s of each-other. The group delay obtained from phantoms with higher gelatine concentrations were generally higher, with phantom iii obtaining the largest value of 277 *μ*s on day 2. All of the group delay values are in the *μ*s range and so small compared to many electrophysiological signal components (for example P100 responses in EEG which arise after 100 ms [5]). Nevertheless, the delays are not constant with frequency and this could introduce distortion into a pre-recorded signal played-out through a phantom. If waveform morphology and the timing of different peaks or signal components is critical to the experiment the gelatine phantom is being used in then the pre-recorded signal should be pre-distorted to correct for this.

**Fig. 7.**
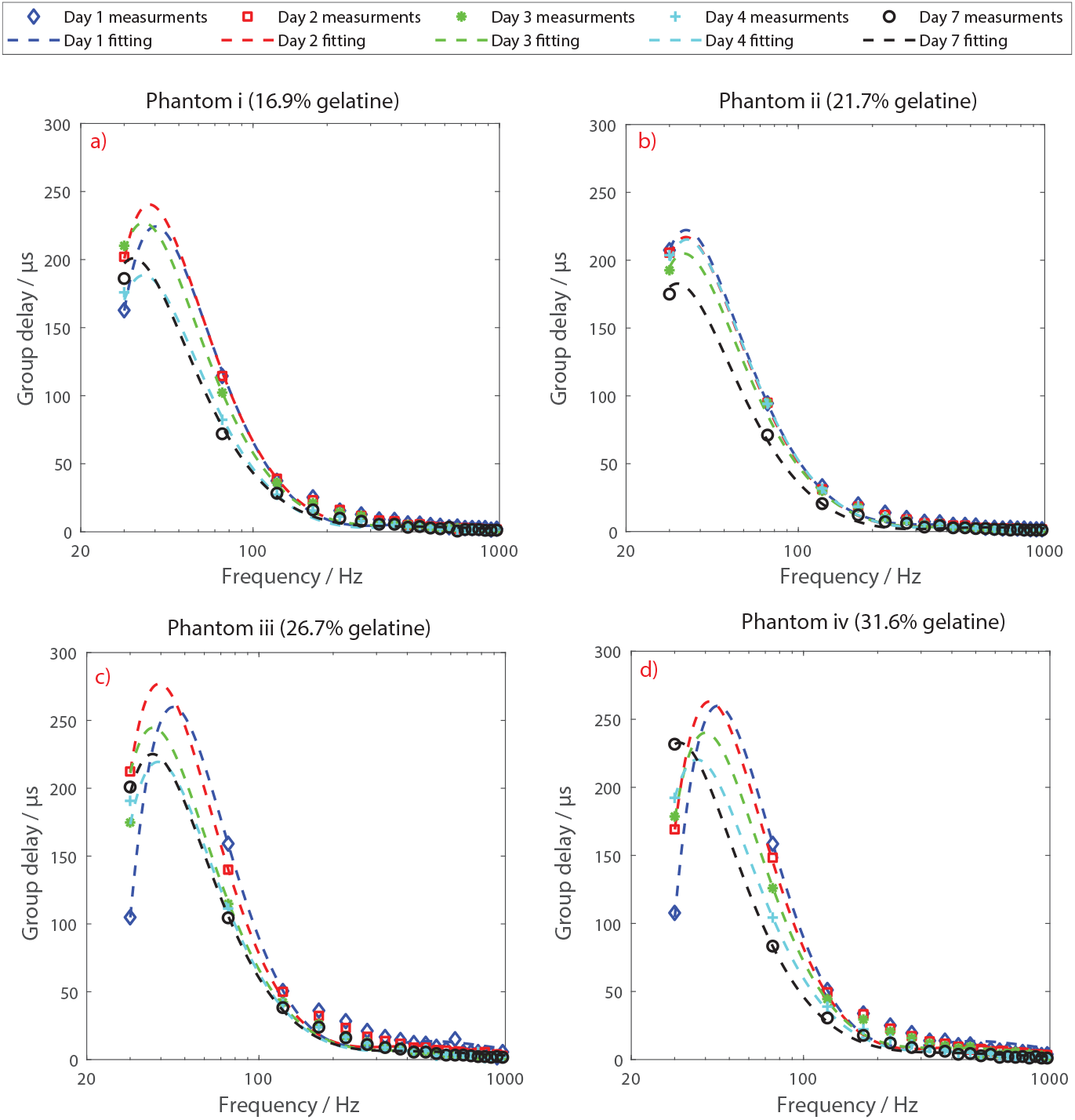
Group delay measurements and fitting lines for phantoms i–iv (in panels a–d respectively).

### B. Effect of NaCl concentration

Fig. 8 shows the contact impedance response (magnitude and phase) of phantoms v–x, having concentrations of NaCl 0–10%, and how this changes over the course of seven days. The measurements show the contact impedance magnitude decreases by a factor of approximately 1.3 for small amounts of added NaCl (0.5–1%) and by a factor of approximately 14 for larger amounts (10% NaCl). This is because the NaCl provides ions for conduction [13] and it allows a wide range of electrical tuning to be performed. This comes at the potential cost of larger phase changes across frequency compared to the NaCl-free phantoms from Fig. 6. Up to −35° for phantom viii with 2% of NaCl is now present. However, this effect is non-linear and increasing the concentration of NaCl further does not lead to larger phase changes. Instead, the phase change across frequencies is less monotonically increasing (as in Fig. 8a-d) with a local decrease in phase appearing at approximately 100 Hz (in Fig. 8f).

**Fig. 8.**
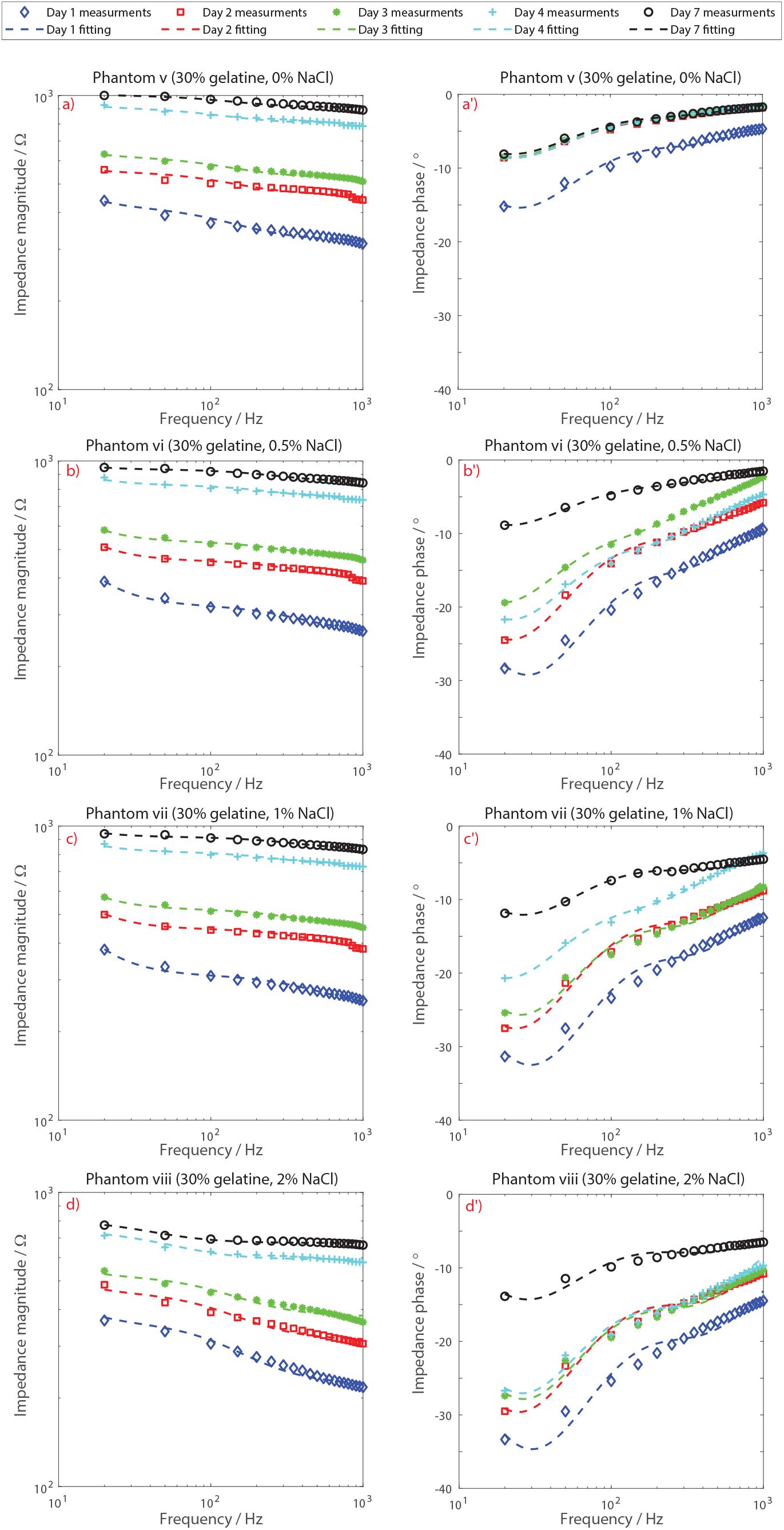
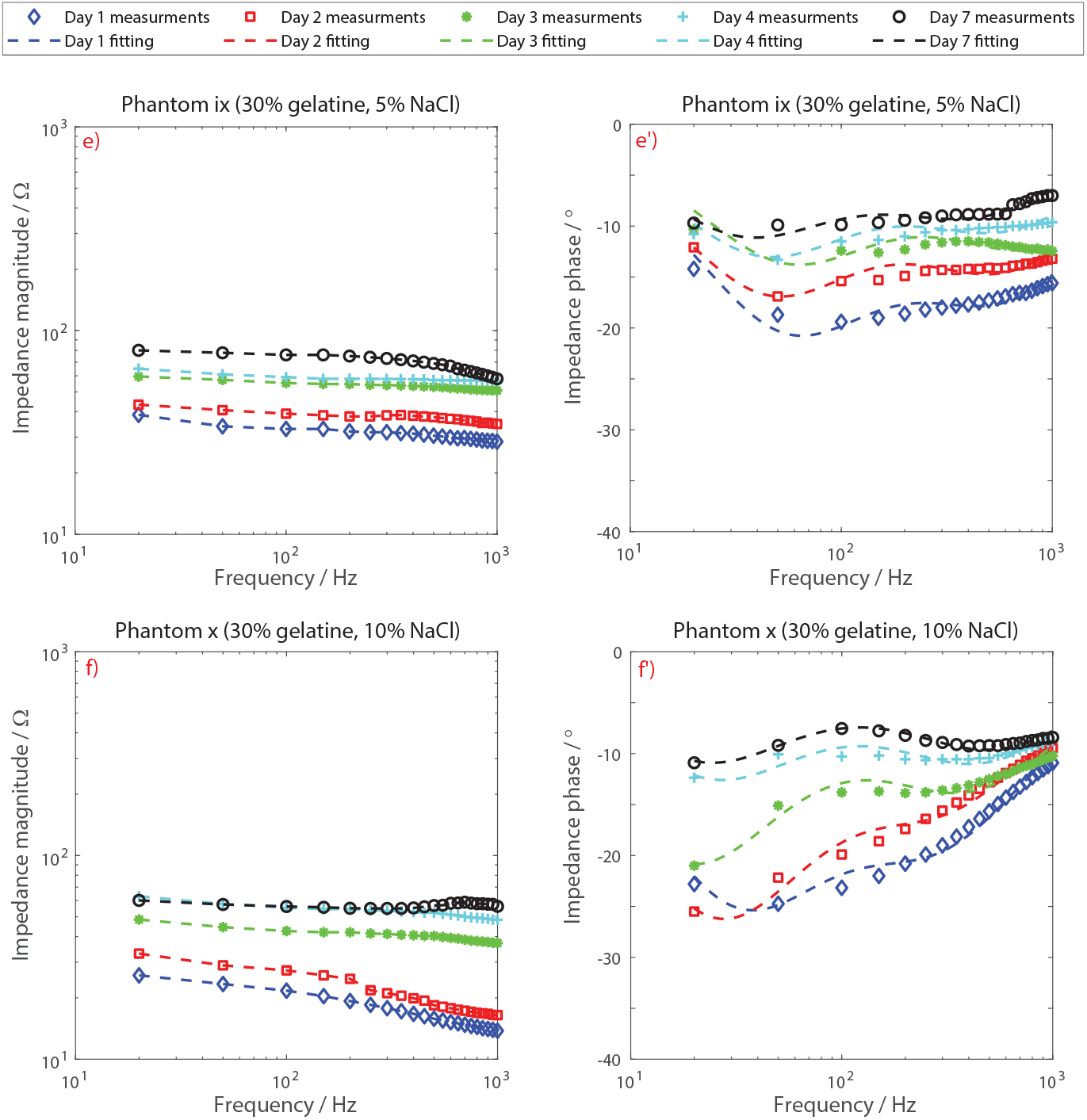
Impedance magnitude and phase responses for gelatine phantoms v–x (in panels a–f respectively) with concentrations of NaCl 0–10%, and how these vary over the course of 7 days. Dashed lines indicate the fit of the second order electrical model from Fig. 2. (Continued on next page.)

The corresponding group delays are shown in Fig. 9. The maximum group delays are substantially increased compared to the NaCl-free phantoms, with up to 558 *μ*s for phantom vi with 0.5% of NaCl. The group delay is also much more variable, particularly at low frequencies (30–100 Hz). Large changes, of up to 400 *μ*s, are present between days. While the addition of NaCl gives greater control over the electrical conductivity and impedance magnitude, this is traded-off with control of the phase properties of the phantom and potentially a decrease in useful life. For many applications signal components being delayed by a few hundred microseconds will not be significant. Nevertheless, if adding NaCl it is much more likely that pre-distortion of any inputted signal will be required if accurate timings of peaks needs to be maintained at the surface of the phantom.

**Fig. 9.**
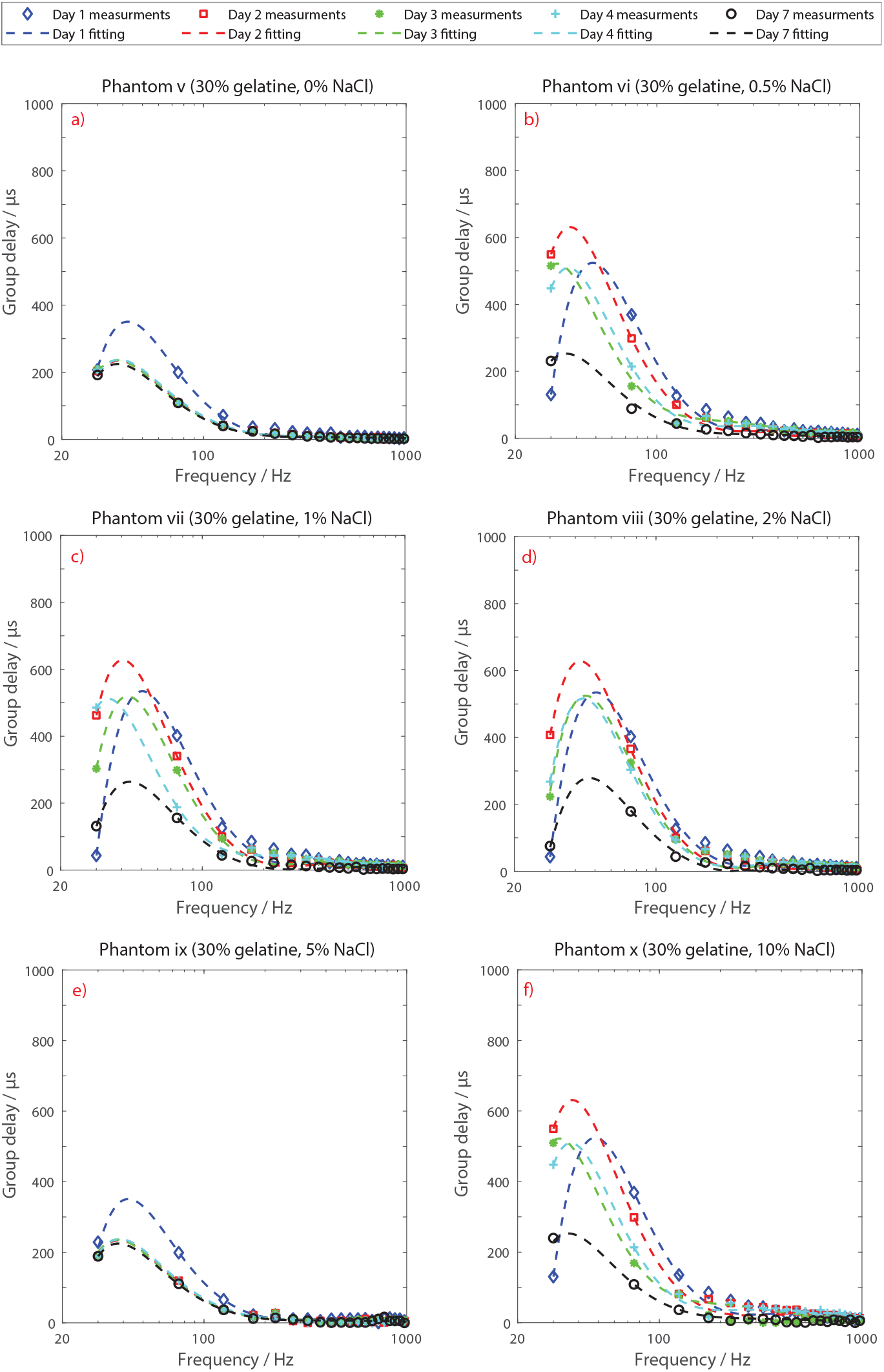
Group delay measurements and fitting lines for phantoms v–x (in panels a–f respectively).

### C. Gelatine phantom repeatability

To assess the repeatability and reusability of the gelatine phantoms 10 copies of phantoms i–iv were made and tested under the same conditions. The phantoms were assessed for a period of one week and the experimental results indicated that the contact impedance and the group delay were very similar across all phantoms. The maximum differences between the original phantoms and the repeated runs in terms of the mean contact impedance magnitude, mean phase, and mean group delay were: 8 Ω, 0.8° and 6.6 *μ*s respectively. These differences are within the measurement accuracy of our impedance analyser and indicate that the gelatine phantoms are systematically repeatable under similar conditions (i.e. methodology, temperature, concentration, and gelatine type). A representative example of one of these many test configurations is shown in Fig. 10 for phantom i. Here the raw measurement points are from the original phantom, while the dashed lines show the fit from a different manufacture of the same phantom configuration.

**Fig. 10.**
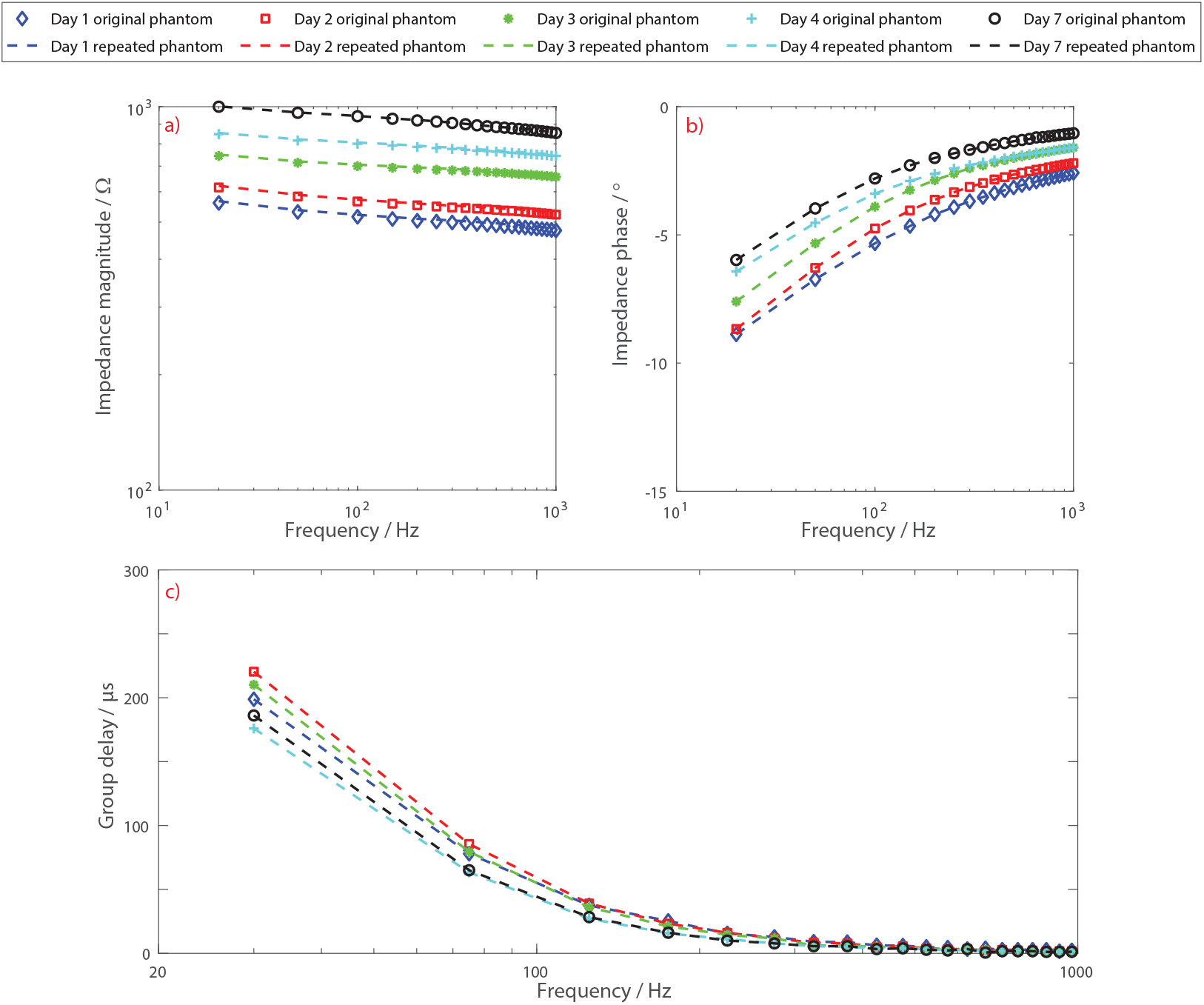
Impedance magnitude, phase, and group delay responses for two manufactured versions of phantom i. Raw measurement points are from the original phantom, while the dashed lines show the fit from a different manufacture of this same phantom configuration and show minimal differences.

### D. Accuracy with porcine skin model

Fig. 11 shows the contact impedance response (magnitude and phase) of a porcine skin sample when assessed on day 1 and day 7 and when compared to the equivalent responses for phantoms i and ii. For the porcine skin the magnitude and phase responses are found to be in the ranges 541–925 Ω and −6.04 – −0.39° respectively, with an average group delay of 12.87–22.43 *μ*s. A comparison between the ex vivo porcine skin model and our gelatine phantoms indicates that phantoms having gelatine concentration of 16.9% and 21.7% (phantoms i and ii) are the closest matches. Table I summarises the ranges of the contact impedance responses and group delays for both porcine skin and phantoms i and ii over the range 20–1000 Hz. This indicates that the electrical properties of the gelatine phantoms are realistic and they fit well within the range of the porcine skin. To our knowledge these results represent the first comparison between the electrical properties of gelatine phantoms and porcine skin and they indicate good proxy between gelatine phantoms and the biological tissue.

**Fig. 11.**
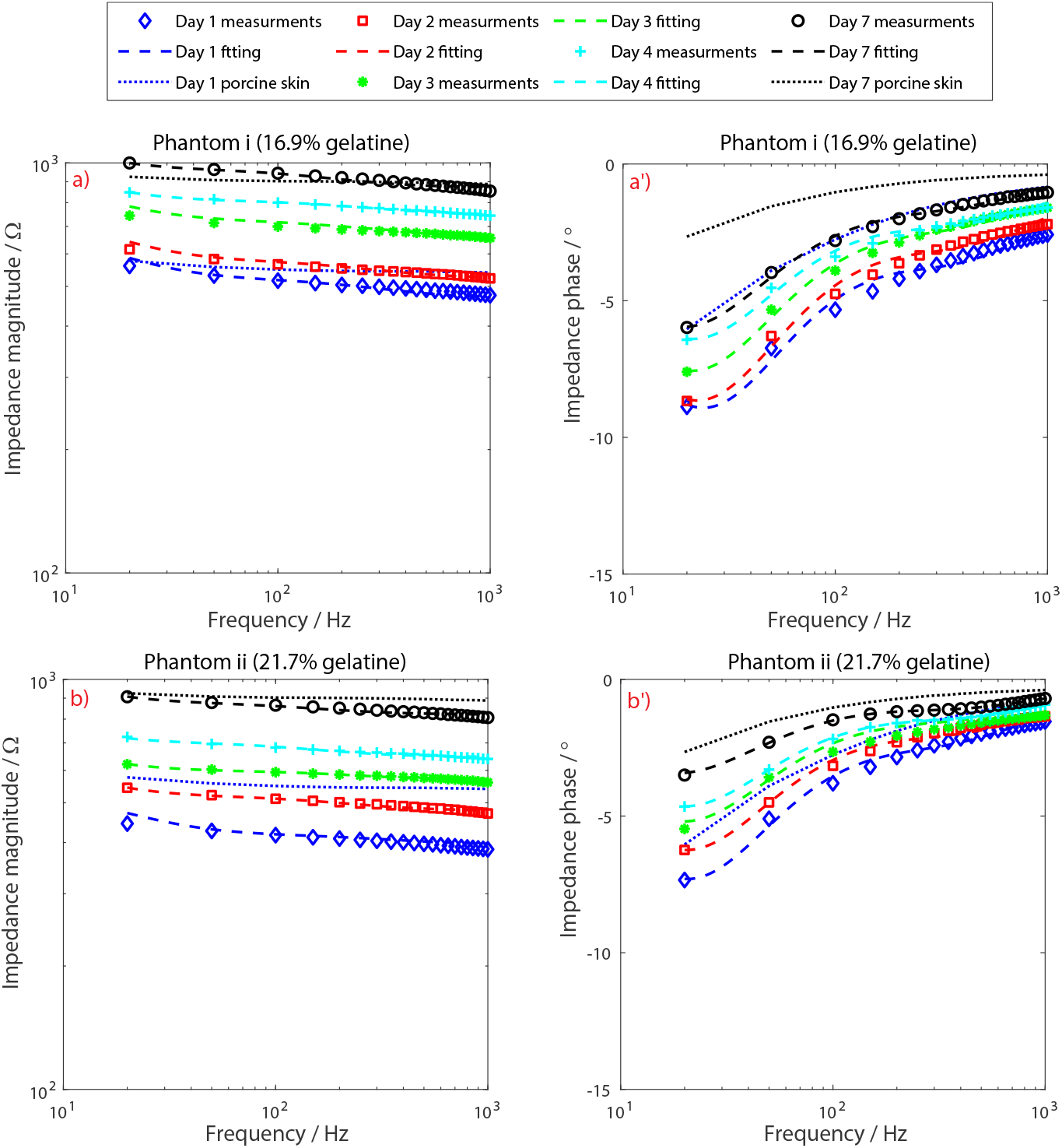
Impedance magnitude and phase responses for a porcine skin compared to the responses from phantom i and ii, compared on day 1 and day 7 with intermediate days plotted for the gelatine phantoms.

**TABLE I.** Comparison of the contact impedance response (given as the max and min values obtained across all frequencies and days) and the average group delay range (averaged across all frequencies obtained across all days) between porcine skin and phantoms i–iv.

| Phantom | Magnitude range / $\Omega$ | Phase range / $^{\circ}$ | Mean group delay / $\mu s$ |
| --- | --- | --- | --- |
| Phantom i | 561 – 854 | –8.88 – –1.04 | 17.0 – 22.4 |
| Phantom ii | 445 – 807 | –7.34 – –0.70 | 15.1 – 20.3 |
| Phantom iii | 302 – 615 | –12.4 – –1.57 | 21.4 – 27.6 |
| Phantom iv | 318 – 606 | –11.7 – –1.06 | 19.8 – 27.3 |
| Ex vivo porcine skin | 541 – 925 | –6.04 – –0.39 | 12.9 – 22.4 |

### E. Accuracy with electrical model

The break-points of the two resistor–capacitor tanks, fitting the collected impedance data to the model from Fig. 2 and which have been plotted on the previous results curves, are given in Table II and Table III for the poles and zeros respectively. These show a good match between the gelatine phantoms, the electrical target model, and the ex vivo porcine skin model. The tunability of the gelatine phantoms is such that the poles and zeros of either the electrical target model or the ex vivo porcine skin model can be matched. Focusing on the poles, particularly for the lower break point, phantom ii and phantom iv match the 6.1 Hz porcine skin model well, with a ~7 Hz cut-off which is also in the range of the electrical target. The higher frequency break-point is more variable, with the mean cut-off of phantom ii (289 Hz) being approximately double the target of the electrical cut-off (159 Hz), and half the porcine skin cut-off (608 Hz). Phantom ii sits in the middle, giving a good trade-off between the different modelling approaches for giving the wanted frequency response. The higher frequency zero resulting from these fits is generally above 1.5 kHz and so above the main frequency range of interest. The lower frequency zero from both phantom ii and the ex vivo porcine skin model is close to 50 Hz and within the range expected to match the electrical target model.

**TABLE II.**
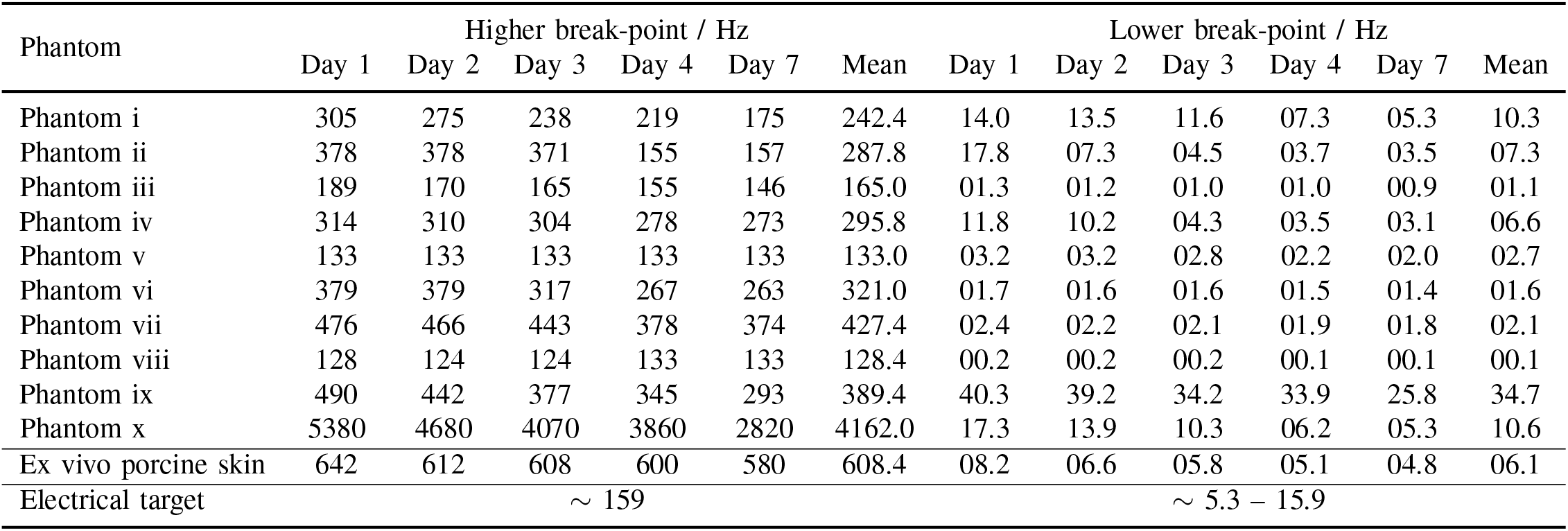
Pole break-points obtained from fitting the electrical model of Fig. 2 to the measured impedances for the 10 different phantom configurations and the ex vivo porcine skin model.

**TABLE III.**
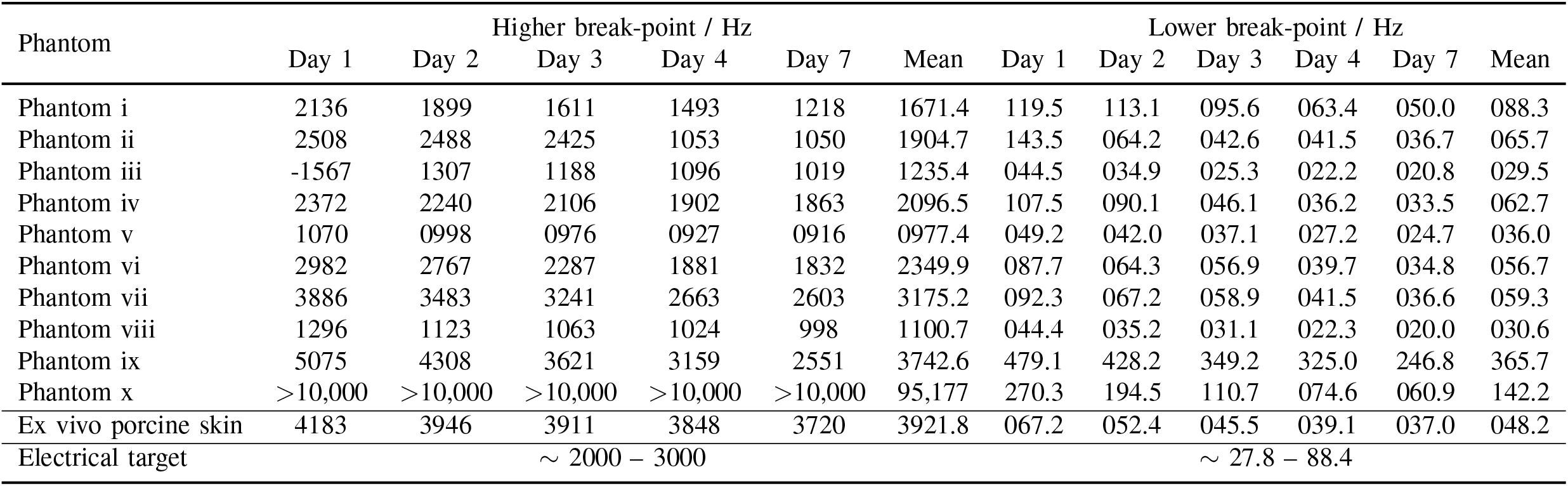
Zero frequencies obtained from fitting the electrical model of Fig. 2 to the measured impedances for the 10 different phantom configurations and the ex vivo porcine skin model.

## VI. Discussion

Gelatine based head phantoms are becoming widely used in the academic literature, but have relatively little public information about their electrical properties (particularly at a.c.) and accuracy. This paper has investigated ten different phantom configurations characterising the impedance path through a cuboid shaped phantom combined with an Ag/AgCl electrode, comparing this to both previously reported electrical models of Ag/AgCl electrodes placed on skin and a model made from ex vivo porcine skin. Further, we have shown how the electrical properties of the phantoms can be tuned using different concentrations of gelatine and of NaCl added to the mixture, and the characterisation has been performed across multiple days of use. The results show that the gelatine based phantoms can accurately mimic the frequency response properties of the body–electrode system to allow for the controlled testing of new electrode designs. The phantoms have also been shown to be repeatable, although this is under the same lab conditions each time (temperature, time of day, no humidity control present), and these conditions may differ from lab-to-lab around the world.

To our knowledge this is the first in-depth characterisation of the a.c. electrical properties of gelatine based phantoms, and the first comparison of the accuracy of the phantoms to both the established double resistor–capacitor tank model of the body–electrode system and to measurements from a biological tissue phantom. While variances are present, all approaches give very similar break-points for the frequency response. Thus, the gelatine phantoms presented in this paper are a cost effective approach for intensive research and can be used to validate measurements under realistic conditions. The phantom allows easy assessment and evaluation of EEG and ECG electrodes and instrumentation, as well as simplifying comparisons in skin to electrode contact impedance between different types of electrodes. The experimental work presented herein provides a reference data set for the contact impedance response of gelatine phantoms and porcine skin and shows variation in this response over time for the first time. This will help in their use as a validation tool where known signals can be passed to novel electrodes to ensure they are collected accurately. Our results will help in tuning the electrical properties of the phantoms using NaCl to match different tissues of interest.

In our characterisation we gave particular attention not only to the impedance of the phantom connection, but also to the group delay present. This group delay was found to increase substantially with the addition of NaCl to the phantom fabrication, going up to several hundred microseconds for some frequencies and phantom configurations. The NaCl based phantoms allowed much greater control over the absolute impedance values present, and in practice we found these phantoms kept for much longer before mould would start to form. However, this potentially comes at the cost of increased signal distortion when they are used to play-out pre-recorded signals as some frequency components will pass through the phantom faster than other signal components, and this changed substantially over time potentially limiting the usable life of the phantom. For phantom tests where the precise signal morphology and the locations of different peaks is important, when adding NaCl it may be necessary to pre-distort the phantom input signal to correct for any group delay effects.

While we found that both gelatine and porcine phantoms gave similar frequency response characteristics, and this was an important accuracy validation step in our work, in principle it means that porcine phantoms could be used in preference to gelatine if desired. While this is the case, our experience leads us to select gelatine phantoms as the preferred approach because it is more manageable, can be generated in the lab at any time, and gives more control over the electrical properties, unlike biological tissue. Both of these models, in our work, have the limitation of no hair being present. In practical electrode applications, particularly for EEG on the head, hair is a major obstacle that lengthens the setup time required (as the hair needs to be parted in order to make contact with the scalp) and this makes getting a good electrical contact difficult. In contrast we obtain a good electrical contact every time. To our knowledge there are no gelatine models to date that have successfully incorporated hair, and this is an important area of future investigation. We also have not attempted to mimic the mechanical properties of the body together with the electrical properties, and future work should look to combine these. ([15] has investigated the mechanical properties of gelatine for making EEG head phantoms.) Our results are also limited by the frequency range of our test equipment, with 20 Hz as the lower cut-off. For future works, particularly for EEG phantoms, investigation of the 0.1–20 Hz range would be highly beneficial.

For improved phantoms in future work, the addition of mechanical artefacts such respiration and balisocardiogram (BCG) effects would be highly beneficial. These are of particular interest for investigating combined EEG+tES using head phantoms, where small movements due to heart beats are known to introduce artefacts into the EEG collection process [46] and such effects are not modelled in our current approaches. Finally, in this work we have focused on single layer phantoms, but multi-layer phantoms are an emerging area of interest. The bulk conductivity of bones is estimated as 0.013 S/m [47], the skull 0.015 S/m [29], scalp 0.43 S/m [47], brain 0.12–0.48 S/m [47], and heart 0.1 S/m [48], all of which could be replicated by our phantom with different levels of NaCl added. A multi-layer phantom, made only from different formulations of gelatine would thus be possible and may lead to more accurate models in the future.

## VII. Conclusions

In this paper 10 gelatine phantom configurations having different concentrations of gelatine and NaCl were generated and characterised. These phantoms were found to be feasible as a cost-effective surrogate conductive material for measuring electrophsyiological responses. Contact impedance and group delay measurements indicated durability in the electrical response of the phantoms over the course of 7 days. Experimental results showed that adding NaCl shifted the magnitude and the phase responses of the contact impedance and allowed tunability and selectively in the phantom’s electrical properties to match a desired impedance profile of realistic tissue (human and animal) in a repeatable and reusable manner. The contact impedance profile for both gelatine phantoms and porcine skin phantoms fit well with each other and are in-line with the results from an electrical model of wet Ag/AgCl electrodes over the frequency band 20–1000 Hz. These findings indicate the potential for using these phantoms for the design of novel electrode and instrumentation systems as they provide a robust testing platform with a known signal to collect allowing controlled device verification.

## References

[1] A. J. Casson, M. Abdulaal, M. Dulabh, et al., “Electroencephalogram,” in Seamless Healthcare Monitoring, T. Tamura and W. Chen, Eds., Cham: Springer, 2018, pp. 45–81.

[2] A. J. Casson, “Wearable EEG and beyond,” Biomed. Eng. Lett., vol. 9, no. 1, pp. 53–71, 2019.

[3] C. Lee and J. Song, “A chopper stabilized current-feedback instrumentation amplifier for EEG acquisition applications,” IEEE Access, vol. 7, no. 1, pp. 11 565–11 569, 2019.

[4] J. Lee, K. Lee, U. Ha, et al., “A 0.8-V 82.9-μW in-ear BCI controller IC with 8.8 PEF EEG instrumentation amplifier and wireless BAN transceiver,” IEEE J. Solid-State Circ., vol. 54, no. 4, pp. 1185–1195, 2019.

[5] A. Velcescu, A. Lindley, C. Cursio, et al., “Flexible 3D-printed EEG electrodes,” Sensors, vol. 19, no. 7, p. 1650, 2019.

[6] G. Li, J. Wu, Y. Xia, et al., “Towards emerging EEG applications: A novel printable flexible Ag/AgCl dry electrode array for robust recording of EEG signals at forehead sites,” J. Neural Eng., vol. 17, no. 2, p. 026 001, 2020.

[7] F. Marini, C. Lee, J. Wagner, et al., “A comparative evaluation of signal quality between a research-grade and a wireless dry-electrode mobile EEG system,” J. Neural Eng., vol. 16, no. 5, p. 054 001, 2019.

[8] Fluke. (2020). Patient monitor simulators, [Online]. Available: https://www.flukebiomedical.com/products/biomedical-test-equipment/patient-monitor-simulators.

[9] D. L. Collins, A. P. Zijdenbos, V. Kollokian, et al., “Design and construction of a realistic digital brain phantom,” IEEE Trans. Med. Imaging, vol. 17, no. 3, pp. 463–468, 1998.

[10] N. C. Rogasch, R. H. Thomson, Z. J. Daskalakis, et al., “Short-latency artifacts associated with concurrent TMS-EEG,” Brain Stim., vol. 6, no. 6, pp. 868–876, 2013.

[11] L. Tomasevic, M. Takemi, and H. R. Siebner, “Synchronizing the transcranial magnetic pulse with electroencephalographic recordings effectively reduces inter-trial variability of the pulse artefact,” PLoS ONE, vol. 12, no. 9, pp. 1–10, 2017.

[12] D. Veniero, M. Bortoletto, and C. Miniussi, “TMS-EEG co-registration: On TMS-induced artifact,” Clin. Neurophysiol., vol. 120, no. 7, pp. 1392–1399, 2009.

[13] W. D. Hairston, G. A. Slipher, and A. B. Yu, “Ballistic gelatin as a putative substrate for EEG phantom devices,” in IEEE EMBC, Orlando, Aug. 2016.

[14] A. Yu and W. D. Hairston. (2019). Open EEG Phantom, [Online]. Available: https://doi.org/10.17605/osf.io/qrka2/.

[15] E.-R. Symeonidou, A. D. Nordin, W. D. Hairston, et al., “Effects of cable sway, electrode surface area, and electrode mass on electroencephalography signal quality during motion,” Sensors, vol. 18, no. 1073, pp. 1–13, 2018.

[16] S. Kohli and A. J. Casson, “Removal of gross artifacts of transcranial alternating current stimulation in simultaneous EEG monitoring,” Sensors, vol. 19, no. 1, p. 190, 2019.

[17] J. Vosskuhl, T. P. Mutanen, T. Neuling, et al., “Signal-space projection suppresses the tACS artifact in EEG recordings,” bioRxiv, vol. 823153, no. DOI: 10.1101/823153, pp. 1–30, 2019.

[18] P. Tallgren, S. Vanhatalo, K. Kaila, et al., “Evaluation of commercially available electrodes and gels for recording of slow EEG potentials,” Clin. Neurophysiol., vol. 116, no. 4, pp. 799–806, 2005.

[19] L. M. Ferrari, U. Ismailov, J.-M. Badier, et al., “Conducting polymer tattoo electrodes in clinical electro- and magneto-encephalography,” npj Flex. Electron., vol. 4, no. 4, pp. 1–9, 2020.

[20] Army Research Laboratory. (2016). Methods for signal validation / EEG “phantom heads”, [Online]. Available: http://openbci.com/forum/index.php?p=/discussion/637/.

[21] S. Kohli and A. J. Casson, “Towards signal processing assisted hardware for continuous in-band electrode impedance monitoring,” in IEEE ISCAS, Baltimore, May 2017.

[22] D. Kim, J. Jeong, S. Jeong, et al., “Validation of computational studies for electrical brain stimulation,” Brain Stim., vol. 8, no. 5, pp. 914–925, 2015.

[23] J. Zhang, B. Yang, H. Li, et al., “A novel 3D-printed head phantom with anatomically realistic geometry and continuously varying skull resistivity distribution for electrical impedance tomography,” Sci. Rep., vol. 7, no. 4608, pp. 1–9, 2017.

[24] J. Dabek, K. Kalogianni, E. Rotgans, et al., “Determination of head conductivity frequency response in vivo with optimized EIT-EEG,” Neuroimage, vol. 127, no. 1, pp. 484–495, 2016.

[25] T. J. Collier, D. B. Kynor, J. Bieszczad, et al., “Creation of a human head phantom for testing of electroencephalography equipment and techniques,” IEEE Trans. Biomed. Eng., vol. 59, no. 9, pp. 2628–2634, 2012.

[26] R. J. Sadleir, F. Neralwala, T. Te, et al., “A controllably anisotropic conductivity or diffusion phantom constructed from isotropic layers,” Ann. Biomed. Eng., vol. 37, no. 12, pp. 522–2531, 2009.

[27] G. P. Mazzara, R. W. Briggs, Z. Wu, et al., “Use of a modified polysaccharide gel in developing a realistic breast phantom for MRI,” Magnetic Resonance Imaging, vol. 14, no. 6, pp. 639–648, 1996.

[28] K. Shmueli, D. L. Thomas, and R. J. Ordidge, “Design construction and evaluation of an anthromorphic head phantom with realistic susceptibility artifacts,” J. Magn. Reson. Imaging, vol. 26, no. 1, pp. 202–207, 2007.

[29] A. Hunold, D. Strohmeier, P. Fiedler, et al., “Head phantoms for electroencephalography and transcranial electric stimulation: A skull material study,” Biomedical Engineering, vol. 63, no. 6, pp. 683–689, 2017.

[30] S. Baillet, J. J. Riera, G. Marin, et al., “Evaluation of inverse methods and head models for EEG source localization using a human skull phantom,” Phys. Med. Biol., vol. 46, no. 77, pp. 77–96, 2001.

[31] R. M. Leahy, J. C. Mosher, M. E. Spencer, et al., “A study of dipole localization accuracy for MEG and EEG using a human skull phantom,” in 4th Int. Conf. on Functional Mapping of the Human Brain, Montreal, Aug. 1998.

[32] M. A. Kandadai, J. L. Raymond, and G. J. Shaw, “Comparison of electrical conductivities of various brain phantom gels: Developing a ‘brain gel model’,” Mater. Sci. Eng. C, vol. 32, no. 8, pp. 2664–2667, 2012.

[33] D. Richler and D. Rittel, “On the testing of the dynamic mechanical properties of soft gelatins,” Exp. Mech., vol. 54, no. 1, pp. 805–815, 2014.

[34] A. I. Farrer, H. Odeen, J. de Bever, et al., “Characterization and evaluation of tissue-mimicking gelatin phantoms for use with MRgFUS,” J. Ther. Ultrasound, vol. 3, no. 9, pp. 1–11, 2015.

[35] Y. M. Chi, T.-P. Jung, and G. Cauwenberghs, “Dry-contact and noncontact biopotential electrodes: Methodological review,” IEEE Rev. Biomed. Eng., vol. 3, no. 1, pp. 106–119, 2010.

[36] J. R. Rice, R. H. Milbrandt, E. L. Madsen, et al., “Anthropomorphic 1 H MRS head phantom,” Med. Phys., vol. 25, no. 7, pp. 1145–1156, 1998.

[37] A. J. Riordan, M. Prokop, M. A. Viergever, et al., “Validation of CT brain perfusion methods using a realistic dynamic head phantom,” Med. Phys., vol. 38, no. 6, pp. 3212–3221, 2011.

[38] K. Miller, K. Chinzei, G. Orssengo, et al., “Mechanical properties of brain tissue in-vivo: Experiment and computer simulation,” J. Biomech., vol. 33, no. 1, pp. 1369–1376, 2000.

[39] Classikool. (2019). 240 Bloom pigskin gelatine professional grade, [Online]. Available: https://www.classikool.com/.

[40] S. Kohli, S. Krachunov and A. J. Casson. (2020). 3D printer molds for head phantom shapes, [Online]. Available: http://dx.doi.org/10.17632/m27cdy3z4c.1.

[41] M. S. Said, N. Seman, and H. Jaafar, “Characterization of human head phantom based on its dielectric properties for wideband microwave imaging application,” Jurnal Teknologi, vol. 73, no. 6, pp. 43–49, 2015.

[42] A. Summerfield, F. Meurens, and M. E. Ricklin, “The immunology of the porcine skin and its value as a model for human skin,” Mol. Immunol., vol. 66, no. 1, pp. 14–21, 2015.

[43] A. Y. Owda, M. Owda, and N.-D. Rezgui, “Synthetic aperture radar imaging for burn wounds diagnostics,” Sensors, vol. 20, no. 3, pp. 1–17, 2020.

[44] A. Y. Owda, N. Salmon, S. Shylo, et al., “Assessment of bandaged burn wounds using porcine skin and millimetric radiometry,” Sensors, vol. 19, no. 13, pp. 1–18, 2019.

[45] M. Ibrani, L. Ahma, and E. Hamiti, The Age-Dependence of Microwave Dielectric Parameters of Biological Tissues. London: InTech, 2012.

[46] N. Noury, J. F. Hipp, and M. Siegel, “Physiological processes non-linearly affect electrophysiological recordings during transcranial electric stimulation,” NeuroImage, vol. 140, no. 1, pp. 99–109, 2016.

[47] T. F. Oostendorp, J. Delbeke, and D. F. Stegeman, “The conductivity of the human skull: Results of in vivo and in vitro measurements,” IEEE Trans. Biomed. Eng., vol. 47, no. 11, pp. 1487–1492, 2000.

[48] S. Gabriel, R. W. Lau, and C. Gabriel, “The dielectric properties of biological tissues: II. measurements in the frequency range 10 Hz to 20 GHz,” Phys. Med. Biol., vol. 41, no. 11, pp. 2251–2269, 1996.

